# Anti-muscle Atrophy Effects of *Lactobacillus reuteri* ATG-F4 and Its Impact on Intestinal Microbiota and Metabolites: Implications for Prophylaxis and Therapy

**DOI:** 10.1101/2023.07.05.546789

**Authors:** Daeyoung Lee, Young-Sil Lee, Gun-Seok Park, Juyi Park, Seung-Hyun Ko, You-Kyung Lee, Do Yeun Jeong, Yong Hyun Lee, Jihee Kang

**Affiliations:** Atogen Co., Ltd., Daejeon, Republic of Korea

**Keywords:** *Lactobacillus*, Probiotics, Muscle atrophy, Inflammation, Gut microbiota, Short chain fatty acids (SCFAs)

## Abstract

*Lactobacillus reuteri* ATG-F4, human gut-derived bacteria, was orally administrated in a model of hindlimb immobilization and confirmed the muscular performance, muscle mass and mechanism on anti-atrophy study. Concomitantly, the changes in the intestinal flora, the metabolites and cytokines were investigated. In the stapled immobilization mice model, ATG- F4 treated group had significantly increased muscle mass, myofiber size, running time to be exhausted and grip strength. The cytokine levels in serum and muscle tissues were reduced by ATG-F4 treatment. Furthermore, the phosphorylation of proteins involved in muscle synthesis such as mTOR, p70S6K, rpS6 and 4E-BP1 increased and MuRF1 related to muscle atrophy factor reduced in the TA and GA muscles. ATG-F4 treatment changed the ratio of main intestinal microflora by increasing the family Muribaculaceae (*phylum Bacteroidetes*) and decreasing the family Lachnospiraceae (*phylum Firmicutes*) and Lactobacillaceae (*phylum Firmicutes*). Also, the level of short chain fatty acids (SCFAs) including butyric acid and acetic acid in the serum of ATG-F4 group were increased. These results suggest that *L. reuteri* ATG- F4 can inhibit muscle atrophy and it is associated with the microbiota and its metabolites with the anti-inflammation effect. ATG-F4 may be a potential prophylactic or therapeutic composition for muscle atrophy.

## 1 Introduction

Probiotics are one of the fastest-growing categories with scientifically proven therapeutic evidence [1, 2]. Some probiotic strains promote the health of your gut microbiome, which can help prevent some common diseases, such as obesity, diabetes, autism, osteoporosis, and some immune disorders [3].

From another point of view, a number of research papers have been reported that probiotics can also affect each other in various organs such as the brain, liver, lungs, bones, and muscles [4-8]. Since probiotics cannot act directly on organs other than the digestive system, the changes of metabolites or immune functions by probiotics may affect various organs. That is, the regulation of intestinal microbes using probiotics is important for improving organ function, which is emerging as an alternative to developing therapeutic agents.

Basically, skeletal muscle is an important tissue that moves the human body, and has functions such as storing carbohydrates as an energy source and regulating body temperature. Meanwhile, muscle mass can be easily lose depending on nutritional status, hormonal balance, physical activity, injury, illness and aging. In particular, muscle atrophy occurs in diseases such as cancer, acquired immunodeficiency syndrome (AIDS), Duchenne muscular dystrophy (DMD), renal and heart failure, chronic obstructive pulmonary disease (COPD), and diabetes [9, 10]. Muscle atrophy adversely affects the recovery and longevity, and deteriorates the quality of life, because muscle mass can influence general metabolism, locomotion and respiration [11]. Therefore, research and development for the prevention or treatment of muscle atrophy are one of the challenges to be solved for human health.

Research papers in associated with relationship between muscle and gut microbiota have been reported over the years [12, 13]. Changes in gut microbiota by prebiotics increased muscle mass in obese and type 2 diabetic mice models [14]. *Akkermansia muciniphila* and *Faecalibacterium prausnitzii* strains inhibited muscle atrophy factor such as Atrogin-1, MuRF1 [15] and *Bifidobacterium breve* B-3 activated the mTOR-p70S6K signaling pathway in muscle [16]. These studies suggest a link between gut microbes and muscle metabolism.

In this study, we utilized the hindlimb immobilization to induce a physical muscle atrophy of mice. There is currently no study investigating the effects of probiotics on gut microbiota and muscle function in a hindlimb immobilization model. Since the hindlimb immobilization is effective in reducing skeletal muscle mass and increasing intramuscular cytokines, it could affect the gut microbiota.

Intestinal dysbiosis affects local and systemic inflammation of the host, and can affect metabolic diseases and specific organs. We hypothesized that intestinal dysbiosis may be associated with muscular atrophy, and that disturbances in the intestinal microbiota and host relationship may exist. In the present study, we investigated the effects of *Lactobacillus reuteri* ATG-F4 on gut microbiota and muscle mass, using a staple-immobilization model [17].

## 2 Methods and materials

### 2.1 Lactobacilli strains

*Lactobacillus reuteri* ATG-F4 strain was isolated from newborn in the previous study [18]. ATG-F4 was cultured on de Man Rogosa Sharpe (MRS) medium (Difco Laboratories, USA) at 37°C for 24h. The cultured ATG-F4 was centrifuged at 4,000 × g for 10 minutes to obtain only the cells, then washed two times by PBS and diluted and then used for animal experiment.

### 2.2 Bioethics declaration

Ethics approval for animal study was provided by the Institutional Animal Care and Use Committee (IACUC) of AtoGen Co., Ltd., and the registration number is ATG-IACUC-RDSP- 200713.

### 2.3 Animals and experimental groups

Five-week-old C57BL/6J male mice were purchased from Central Lab. Animal Inc. (Seoul, Korea). The facility was maintained at a temperature of 23 ± 2 ℃ and a humidity of 55 ± 10 %, with a 12-h light/dark cycle. Mice fed a normal diet (Cargill Inc, Purina®, Korea) and drank sterilized distilled water. Mice were randomly divided into three treatment groups: Untreated (n = 10), Stapled (n = 10), Stapled + ATG-F4 (n = 10) after a 1-week acclimation period.

ATG-F4 was administered orally at a dose of approximately 4.0 × 10^9^ CFU/mice for a total of 27 days, while the untreated group and staple fixation group were administered PBS. On day 14 of the treatment, the right leg of each mouse in the staple fixation group was fixed with a staple for 10 days. Three days after staple removal, fresh fecal samples were collected from each mouse in an empty cage. Additionally, a wire hanging test and a treadmill test were performed on all animals to assess the improvements in muscle rehabilitation and exercise capacity due to ATG-F4 consumption.

All animals were sacrified the day after muscular performance test. Blood samples were collected after euthanasia by CO2 inhalation, and freshly obtained serum samples were processed for cytokines and metabolome analysis. Five types of leg skeletal muscle such as tibialis anterior (TA), gastrocnemius (GA), plantaris (PL), extensor digitorum longus (EDL) and soleus (SOL) were separated and their weights were compared. All muscle tissues were stocked in liquid nitrogen and moved to -70 ℃ until they were analyzed.

### 2.4 Wire hang test

The wire hang test was performed using a 440×330 mm square of wire mesh consisting of 10 mm each in width and length per diameter wire [19]. The time was measured after the mouse was placed to the center of the wire screen and turned over. The screen was fixed at least 40 cm up above a padded surface to absorb the shock when it fell. All animals had warming-up for three minutes on the wire without a weight. The experiment was conducted by hanging a weight of approximately 25% of mice body weight and maximum time set 300 seconds. The wire hang test was repeated three times independently. Recording the falling time of the mouse was compared and analyzed by the average value. Furthermore, the falling time was re-evaluated as score following as; falling time between 1-60 seconds = 1, falling time between 61-120 seconds = 2, falling time between 121-180 seconds =3, falling time between 181-240 seconds =4, falling time between 241-300 seconds =5, falling time after 300 seconds =6.

### 2.5 Treadmill test

The treadmill test was performed with reference to the protocol used for the running test using 5 LANE TREADMILL for MICE (Harvard Apparatus, USA) [20, 21]. Before starting the experiment, the mice were warmed up at a speed of 8 m/min for five minutes. The starting point was 1.25 mA of electrical stimulation to induce mouse running. For the actual test, starting at a speed of 10 m/min, settings were made to automatically increase the speed by 2 m/min every 10 minutes. The running time was measured up to 80 minutes in eight steps. After half hour, the angle was started to increase to two degrees every 10 minutes, the angle was increased up to 10°. Despite the application of electrical stimulation, the experiment was terminated on assumption that the mice were exhausted when mice got electrical shock five times at one step. After the experiment was completed, the measured running time and distance of the mouse were recorded to analyze.

### 2.6 Cytokine assay

The levels of TNF-α and IL-6 in the serum and the supernatant of the crushed muscle tissues were measured using a commercial kit (mouse TNF-α and IL-6 ELISA max standard set, BioLegend). The values of TA and GA muscles were revised to the protein concentration measured by BCA kit and expressed as pg/mg.

### 2.7 Western blot analysis

Flash-frozen TA and GA muscles were thoroughly crushed in liquid nitrogen. And RIPA buffer (0.5M Tris-HCl, pH 7.4, 1.5M NaCl, 2.5% deoxycholic acid, 10% NP-40) containing a protease inhibitor cocktail (Millipore, USA) was added to the crushed muscle tissues after liquid nitrogen was evaporated. The homogenate was centrifuged at 14,000 rpm for 10 min at 4°C, and the supernatant was recovered. Protein concentrations in the supernatant were determined using BCA assay (Thermo Fisher, USA). The protein extract (40μg) was separated on 8 or 10% polyacrylamide mini gel and transferred to a PVDF membrane (Bio-Rad, USA). Blotted membranes were incubated overnight at 4°C in SuperBlock(PBS) Blacking buffer (pH 7.4) containing Kathon^TM^ Anti-microbial Agent with anti-mTOR (1:1000; Cell signaling), anti-p-mTOR (1:1000; Cell signaling), anti-p70S6K (1:1000; Cell signaling), anti-p-p70S6K (1:1000; Cell signaling), anti-rpS6 (1:1000; Cell signaling), anti-p-rpS6 (1:1000; Cell signaling), beta actin (1:1000; Cell signaling). The blots were washed four times with 0.1% tween TBS before being incubated for 1 h at room temperature with goat Anti-rabbit IgG HRP conjugated secondary antibody (Bio-Rad, USA) in 0.1% tween TBS buffer containing 3% BSA (Bovogen, USA). After extensive washes with 0.1% tween TBS, the immunostained bands were revealed with ECL (Bio-Rad, USA). the target complex was detected by ChemiDoc^TM^ Imaging System (Bio-Rad, USA). The target band intensity was quantified using the Image Lab^TM^ software (Bio-Rad, USA).

### 2.8 Real-time PCR

Total RNA was extracted from flash-frozen TA and GA muscles using TRIzol (Thermo Fisher, USA) after crushed them in liquid nitrogen. The homogenate keeping on ice was added chloroform and centrifugated 13,000g, 4℃ for 5min and the supernatant was recovered. Isopropanol which was set 70% of final concentration were added to get RNA precipitate from the homogenate by centrifugation (13,000g, 4℃ for 5min), and this performance was repeated twice by 70 % of ethanol. DEPC water was added to RNA precipitation after drying solvent at room temperature. cDNA was synthesized from the 1 μg of RNA using AccPower RT PreMix (Bioneer, Korea). Primers for PCR amplification were as follows: 5’ - ACACTGGTGCAGAGAGTCGG-3’ (forward) and 5’- TAAGCACACAGGCAGGT CGG-3’ (reverse) for Atrogin-1 and 5’-GCGTGACCACAGAGGGTAAAGA-3’ (forward) and 5’ - GTGGGGAGCCCTATGCTAGTC-3’ (reverse) for MuRF1. Beta actin (GenBank accession no. NM_007393) was amplified by PCR from AccuPower qPCR primer A set (Bioneer, Korea). The quantitative RT PCR assays were performed using 10 ng of cDNA with PowerUP^TM^ SYBR^TM^ Green Master Mix (Thermo Fisher Scientific, USA) and primers by 7500 Fast real-time PCR System (Thermo Fisher Scientific, USA). qPCR conditions for all reactions included an initial 20 seconds denaturation step at 95℃, followed by 45 cycling stages 3 seconds at 95℃ and 30 seconds at 60℃.

### 2.9 Histology

TA and GA muscles were excised, fixed in 10% formalin for more than 48h at room temperature. The fixed tissue was subjected to general tissue processing procedures such as trimming, dehydration, paraffin embedding, and thinning, to prepare a specimen for histopathological examination, and then staining Hematoxylin & Eosin (H&E). Histopathological changes were observed by H&E slide pictures which were taken a 400-fold enlargement using an optical microscope (Olympus BX53, Japan). Myofiber size results are obtained from the cross-sectional area (CSA) of H&E stain sections. Over 30 myofibers/field from 3 different views was examined.

### 2.10 Fecal microbiota analysis

Genomic DNA extraction from fecal samples was performed using the QIAamp PowerFecal Pro DNA Kit (Qiagen, Germany). The quantity and quality of extracted DNA were measured using Qubit 3.0 Fluorometer (Thermo Fisher Scientific, USA) and agarose gel electrophoresis, respectively. The V4 hypervariable regions of the bacterial 16S rRNA were amplified with unique 8 bp barcodes and sequenced on the Illumina iSeq 100 system plATGorm according to standard protocol [22]. The raw sequence data were submitted to the NCBI’s SRA database (NCBI BioProject, raw data accession number: PRJNA694467). Raw reads were analyzed using the QIIME2 pipeline [23]. Sequences were quality filtered and clustered into operational taxonomic units at 97% sequence identity according to the SILVA 132 database [24]. The operational taxonomy units were identified at phylum to family levels.

### 2.11 Short chain fatty acids analysis

Acetic acid (≥99.8% purity), butyric acid (≥99.5% purity) and propionic acid (≥99.5% purity) were of analytical grade purchased from Sigma-Aldrich (St. Louis, MO, USA). Ammonium formate (≥99.995% purity), formic acid (≥99.5% purity), high-performance liquid chromatography (HPLC) grade methanol (MeOH), HPLC grade acetonitrile (ACN) and HPLC grade water were purchased from Sigma-Aldrich. A 100 μL aliquot of serum was mixed with 500 μL of ice-cold MeOH with 1 % formic acid. The solution was mixed by vortex for 20 minutes and left for 2 hours (4°C) to solidify the protein precipitate. After centrifugation at 14,000 rpm for 30 min at 4◦C, the supernatant was filtrated by using a PVDF syringe filter (0.22 μm pore size) (Millipore, Billerica, MA). The filtrated sample was injected into the liquid chromatography-tandem mass spectrometry (LC-MS/MS) system. LC-MS/MS analyses were carried out using an Exion LC system connected to a QTRAP 4500 mass spectrometer (AB SCIEX, Framingham, MA, USA). The LC analyses were carried out using an Intrada Organic Acid column (150 × 2 mm, 3-μm particle size, Imtakt Kyoto, JPN). A 5 μL aliquot was used for the autosampler injection and the flow rate was 0.2 mL/min at 40 °C. The negative ion mode scanning of a gradient mobile phase, consisting of (A) acetonitrile/water/formic acid (10/90/0.1, v/v) solution and (B) acetonitrile/100 mM ammonium formate (10/90, v/v) solution, was used. The gradient started at 0% B and was held at this value for 1 min. The gradient increased linearly to 100% B in 6 min. The mobile phase composition was held at 100% B for 3 min before it returned to 0% B in 0.1 min. Finally, the gradient was kept at 0% B for 4.9 min to re-equilibrate the column. The total analysis time was 15 min. The analysis was performed using an electrospray ionization source in negative mode. The operation conditions were as follows: ion spray voltage, 4500 V; curtain gas (CUR), 25 psi; collision gas (CAD), medium; ion source gas 1 (GS1) and ion source gas 2 (GS2), 50 and 50 psi; the turbo spray temperature (TEM), 450 °C; entrance potential (EP), 10 V; collision cell exit potential (CXP), 5 V. Nitrogen was used in all cases. Analytes were quantificated by multiple reaction monitoring (MRM) employing the following precursor to product ion transitions and parameters: acetic acid, m/z 105.0 → 44.9 with DP 5 V and CE 18 eV; butyric acid, m/z 133.0 → 44.9 with DP 5 V and CE 18 eV; propionic acid, m/z 119.0 → 45.1 with DP 5 V and CE 16 eV. SCIEX OS 2.0.0 software was used for data acquisition and processing, and Analyst 3.3 software was used for data analysis.

### 2.12 Statistical analyses

All data are expressed as means ± SEM. One-way analysis of variance (ANOVA) with Dunnett’s test was used for multiple comparison by GraphPad Prism 8.0 software. Differences were considered statistically significant at *P* < 0.05. For the fecal microbiota analysis, the non-parametric Kruskal-Wallis test was used to compare the differences in diversity indexes and microbial taxa.

## 3 Results

### 3.1 Effects of *L. reuteri* ATG-F4 on body weight & muscles weight

The total body weights among all three groups showed no significant differences after 10 days of immobilization and three days of staple removal (Figure 1A). However, the muscle mass of TA, GA, PL, EDL, and SOL in the stapled group was atrophied by 28.9%, 30.3%, 28.0%, 20.7%, and 15.3%, respectively, compared to the untreated group. Treatment with ATG-F4 resulted in a significant increase in muscle mass of TA, GA, and PL by 17.3%, 30.3%, and 27.0%, respectively, compared to the stapled group (Figure 1B-G). Additionally, the total muscle mass of the hindlimb was significantly augmented by 22.4% with ATG-F4 compared to the stapled group (Figure 1H).

**Figure 1.**
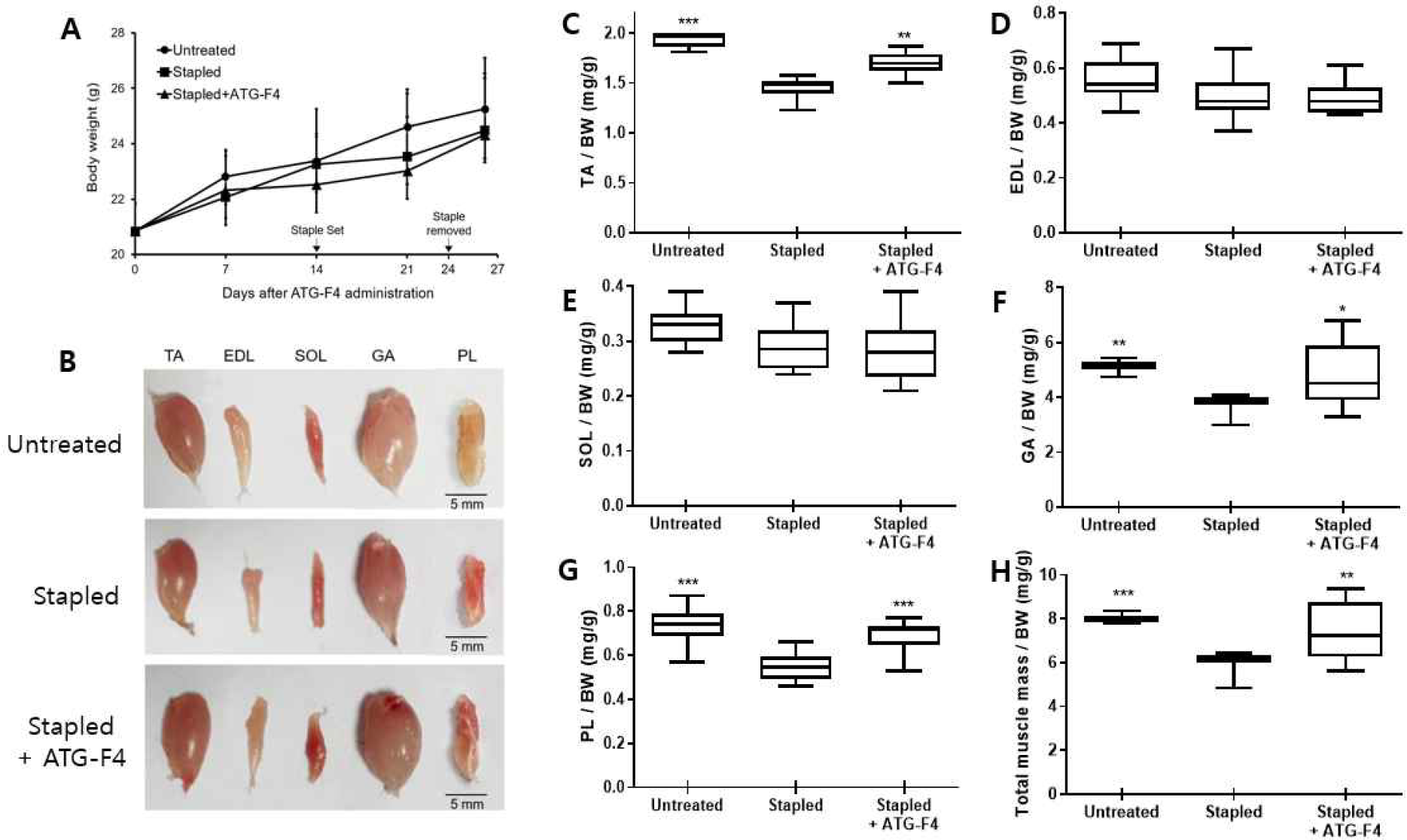
Changes of bodyweight and muscle mass of hindlimb. (A) Bodyweight; (B) Pictures of hindlimb; (C-H) Relative weight of TA (C), EDL (D), SOL (H), GA (F), PL (G) and Total muscle (H); TA: tibialis anterior; EDL: extensor digitorum longus; SOL: soleus; GA: gastrocnemius; PL: plantaris. Untreated: unstapled group; Stapled: hindlimb immobilization group; Stapled + ATG-F4: hindlimb immobilization + *L. reuteri* ATG-F4 (4.0×10^9^ CFU/day) treated group. Data are presented as means ± SEM *p < 0.05, **p < 0.01 and ***p < 0.001 compared to Stapled.

### 3.2 Effects of *L. reuteri* ATG-F4 on myofiber size

we analyzed the cross-sectional area (CSA) of myofibers in the TA and GA muscles of each experimental group, following H&E staining. The average size of myofibers in the TA and GA muscles of the stapled group was decreased compared to the untreated group, while the size of myofibers in both TA and GA muscles of the ATG-F4 group was significantly increased compared to the stapled group (Figure 2A-C). Moreover, there was a significant increase in the percentage of large myofibers in the TA and GA muscles of the ATG-F4 group compared to the stapled group (Figure 2D and E).

**Figure 2.**
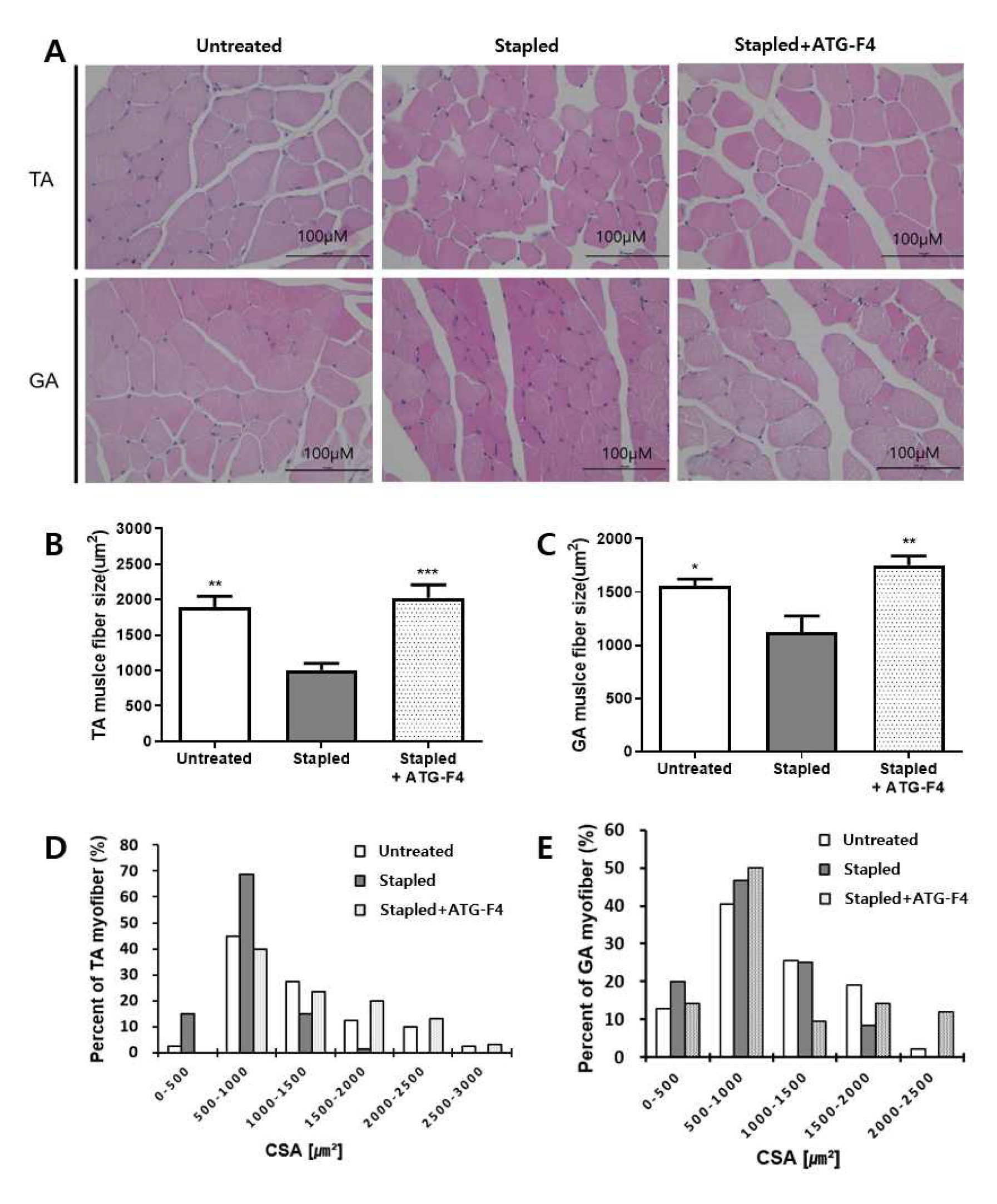
H&E staining of TA and GA muscle fibers. (A) Representative pictures of TA and GA muscle fiber; (B-C) Quantification of CSA in TA (B) and GA (C) muscle fibers; (D-E) Myofiber size distribution of TA(D) and GA(E) muscle. TA: tibialis anterior; GA: gastrocnemius; Untreated: unstapled group; Stapled: hindlimb immobilization group; Stapled + ATG-F4: hindlimb immobilization + *L. reuteri* ATG-F4 (4.0×10^9^ CFU/day) treated group. Data are presented as means ± SEM *p < 0.05, **p < 0.01 and ***p < 0.001 compared to Stapled.

### 3.3 Effects of *L. reuteri* ATG-F4 on muscle function

To estimate muscle strength, we performed the wire hang test 3 days after staple removal, with timing beginning as soon as a mouse faced down on the wire. The latency to fall and average maximum score of ATG-F4-treated mice were significantly higher than those of the stapled group, at 188.9 ± 80.2 vs. 116.1 ± 29.2 and 4.0 ± 1.7 vs. 2.7 ± 0.7, respectively (Figure 3A and B). In the treadmill test to assess muscle functional recovery, the running time to exhaustion for the stapled group mice was significantly shorter than that for the untreated group. In contrast, the ATG-F4 group showed a significant increase in both running time and distance compared to the stapled group, at 48.7 ± 14.2 vs. 34.9 ± 11.2 and 716.6 ± 294.5 vs. 445.9 ± 182.1, respectively (Figure 3C and D).

**Figure 3.**
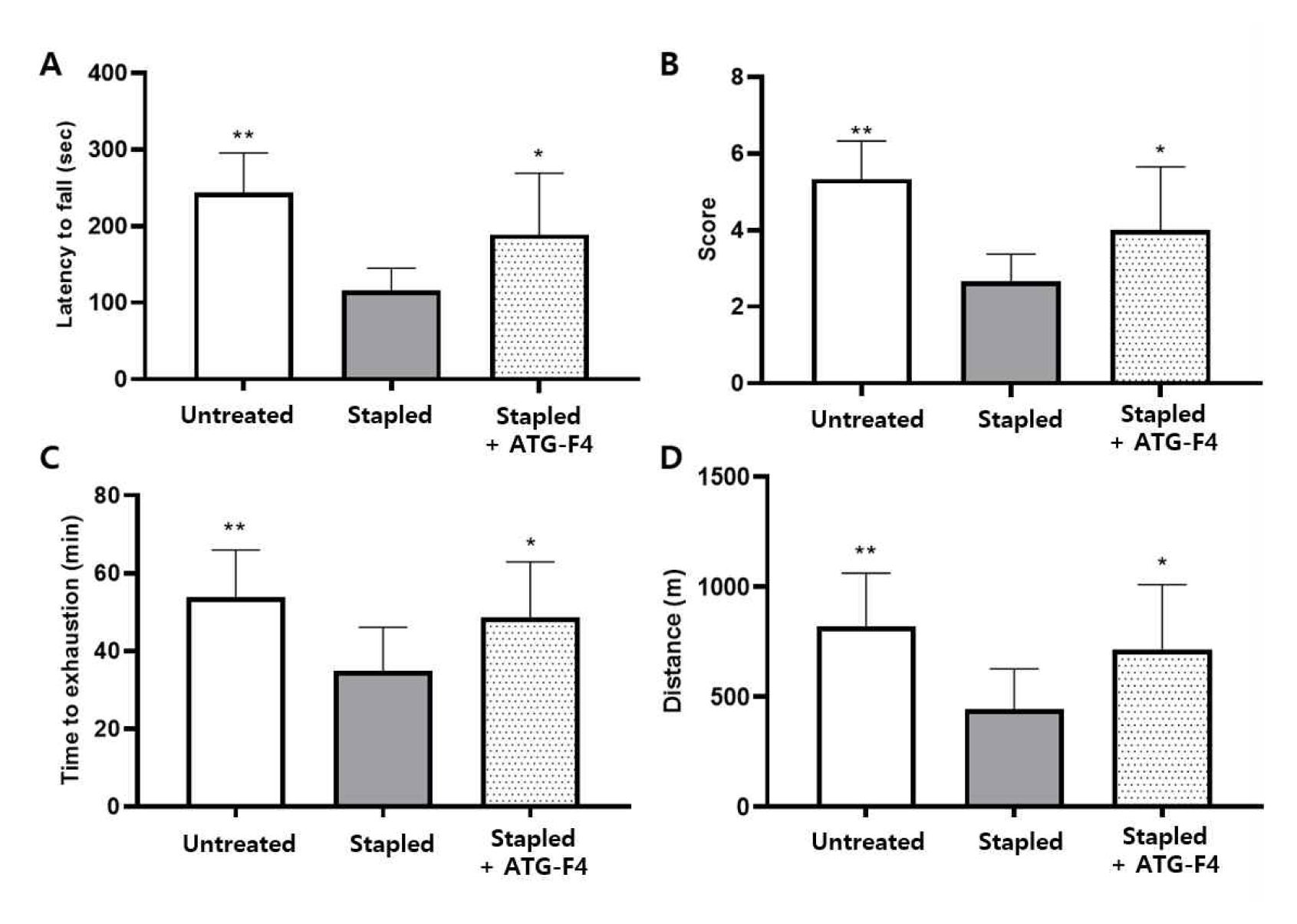
Physical performance of muscle strength and endurance. (A, B) Latency to fall (A) and its score (B) in wire hang test for muscle strength; (C, D) Running time to exhaustion (C) and its distance (D) in treadmill test; Untreated: unstapled group; Stapled: hindlimb immobilization group; Stapled + ATG-F4: hindlimb immobilization + *L. reuteri* ATG-F4 (4.0×10^9^ CFU/day) treated group. Data are presented as means ± SEM *p < 0.05, **p < 0.01 and ***p < 0.001 compared to Stapled.

### 3.4 Effects of *L. reuteri* ATG-F4 on cytokines levels in serum and muscle tissues

The levels of inflammatory cytokines were measured in the serum, TA, and GA muscles. TNF- α and IL-6 levels were increased in the serum and muscle tissues of the stapled group compared to the untreated group. Treatment with ATG-F4 resulted in reduced levels of TNF-α in the serum and TA muscle compared to the stapled group (Figure 4A and B). The level of IL-6 in the serum of the ATG-F4 group was significantly lower than the stapled group (Figure 4D). Moreover, ATG-F4 treatment also decreased the level of IL-6 in TA and GA muscles (Figure 4E and F).

**Figure 4.**
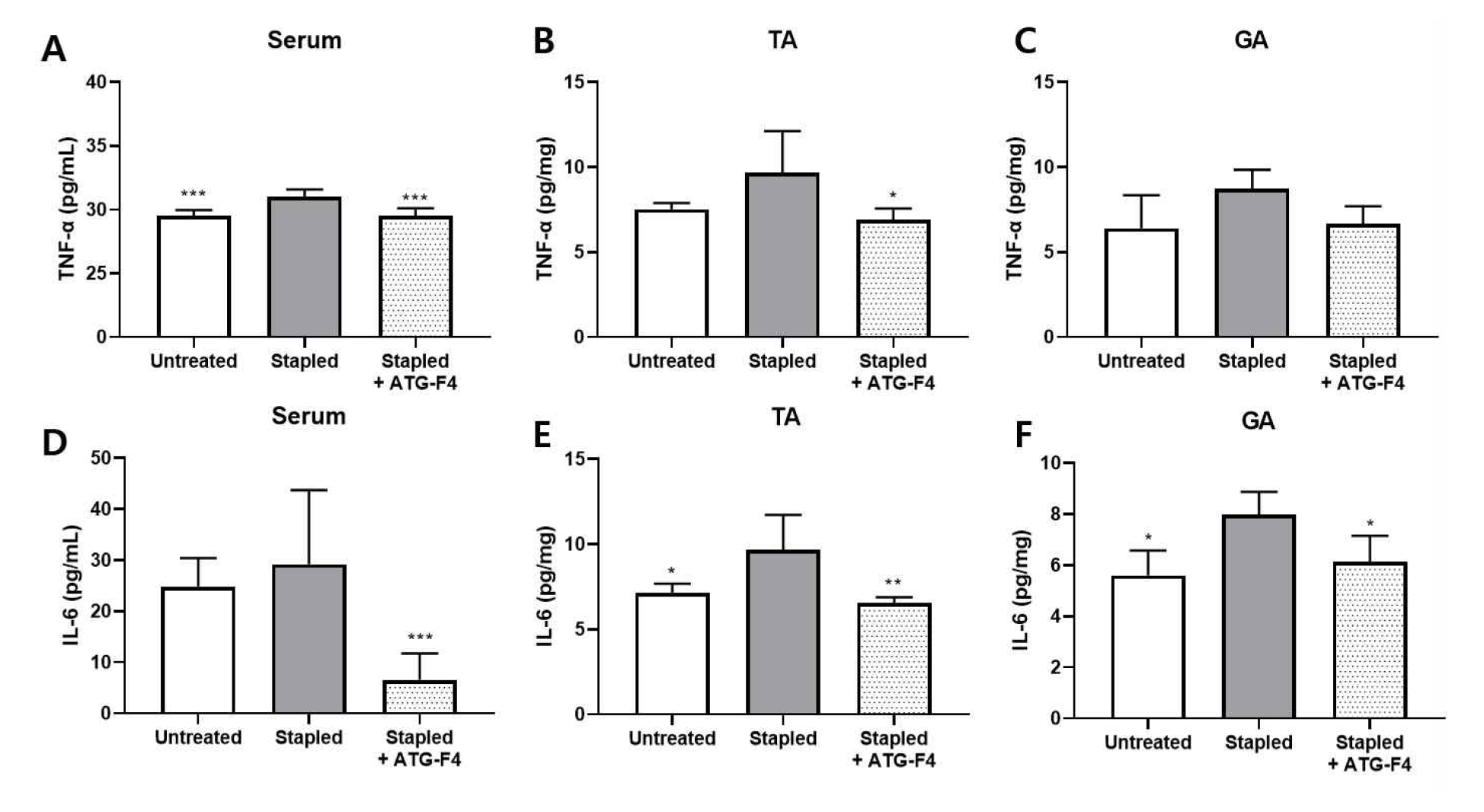
Levels of inflammatory cytokine the serum and muscle tissues. (A-C) The level of TNF-α in serum (A), TA (B), GA (C) muscle; (D-F) The level of IL-6 in serum (D), TA (E), GA (F) muscle of the level of IL-6; TA: tibialis anterior; GA: gastrocnemius. Untreated: unstapled group; Stapled: hindlimb immobilization group; Stapled + ATG-F4: hindlimb immobilization + *L. reuteri* ATG-F4 (4.0×10^9^ CFU/day) treated group. Data are presented as means ± SEM *p < 0.05, **p < 0.01 and ***p < 0.001 compared to Stapled.

### 3.5 Effects of *L. reuteri* ATG-F4 on the expression of muscle atrophy related genes and proteins

The mRNA and protein expression levels of muscle atrophy-related factors such as Atrogin-1 and MuRF1 in the TA and GA muscles were investigated using RT-qPCR and western blot analysis, respectively. In the TA muscle, the expression levels of Atrogin-1 in the stapled group were decreased compared to the untreated group. However, there was no significant difference in the expression level of Atrogin-1 between the ATG-F4 and stapled groups (Figure 5A and C). The expression levels of MuRF1 in the stapled group were not decreased compared to the untreated group, but the expression level of MuRF1 was significantly decreased in the ATG- F4 group compared to the stapled group (Figure 5B and D).

**Figure 5.**
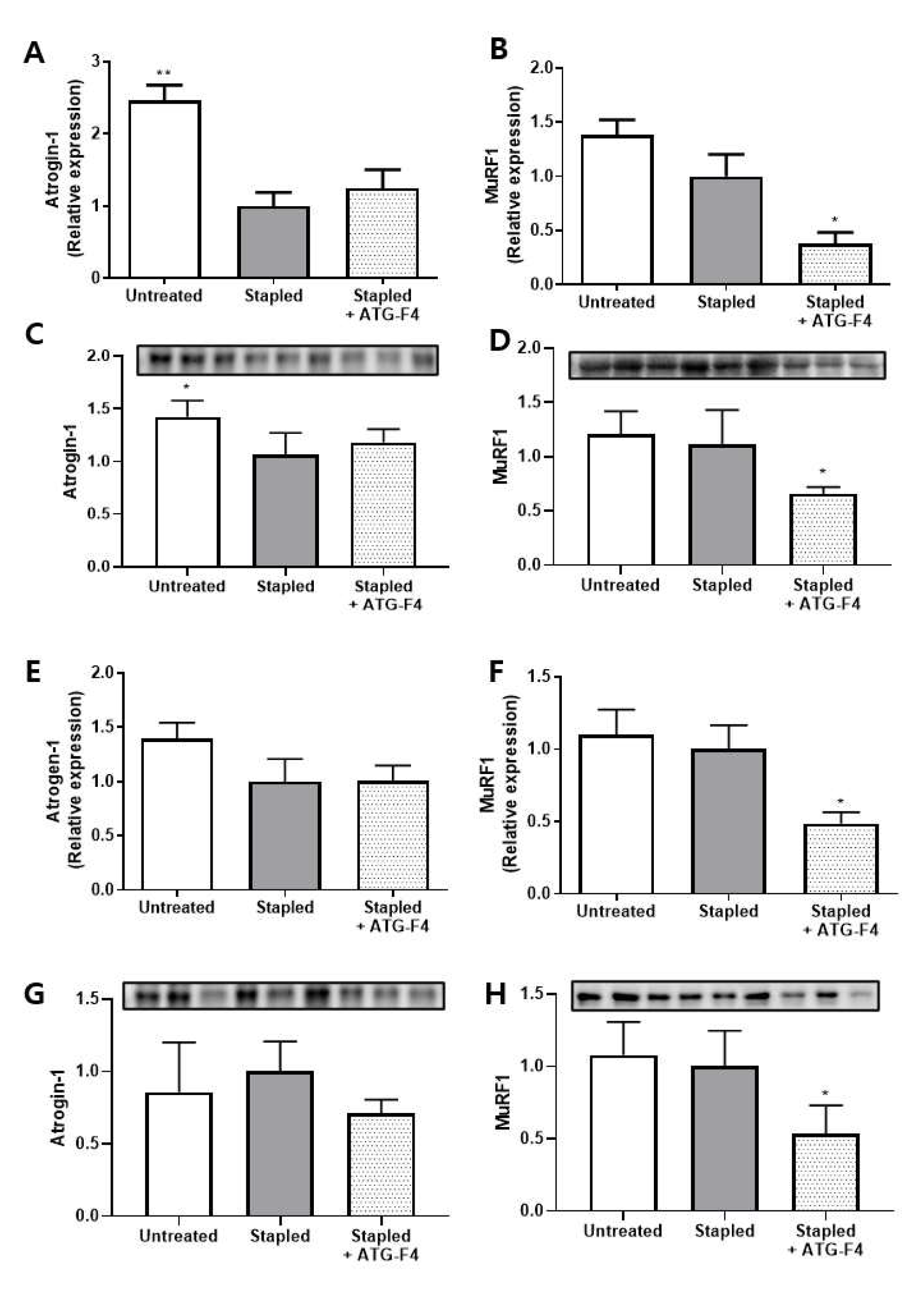
Expression of muscle atrophy factors in muscles. (A, B) The mRNA expression of Atrogin-1 (A) and MuRF1 (B) in TA muscle; (C, D) The protein expression of Atrogin-1 (C) and MuRF1 (D) in TA muscle; (E, F) The mRNA expression of Atrogin-1 (E) and MuRF1 (F) in GA muscle; (G, H) The protein expression of Atrogin-1 (G) and MuRF1 (H) in GA muscle; TA: tibialis anterior; GA: gastrocnemius. Untreated: unstapled group; Stapled: hindlimb immobilization group; Stapled + ATG-F4: hindlimb immobilization + *L. reuteri* ATG-F4 (4.0×10^9^ CFU/day) treated group. Data are presented as means ± SEM *p < 0.05, **p < 0.01 compared to Stapled.

In the GA muscle, the expression levels of Atrogin-1 and MuRF1 in the stapled group were not significantly different from the untreated group (Figure 5E-H). The expression level of Atrogin-1 was not significantly different between the ATG-F4 and stapled groups, however, the expression level of MuRF1 was significantly decreased in the ATG-F4 group compared to the stapled group (Figure 5F and H).

### 3.6 Effects of L. reuteri ATG-F4 on the phosphorylation of proteins related to muscle synthesis in muscles

We investigated the effects of ATG-F4 treatment on the phosphorylation of muscle synthesis-related proteins in TA and GA muscles using the western blotting method. The phosphorylation levels of muscle synthesis-related factors such as mTOR, P70S6K, rpS6, and 4E-BP1 in the TA muscle of the stapled group were not significantly different from those of the untreated group. However, the relative phosphorylation of these proteins in the TA muscle was significantly increased by ATG-F4 treatment (Figure 6A-E). In the GA muscle, the expression levels of muscle synthesis-related factors in the stapled group were either not different or decreased compared to the untreated group. As with the TA muscle, the relative phosphorylation of muscle synthesis factors in the GA muscle was also increased by ATG-F4 treatment (Figure 6F-J).

**Figure 6.**
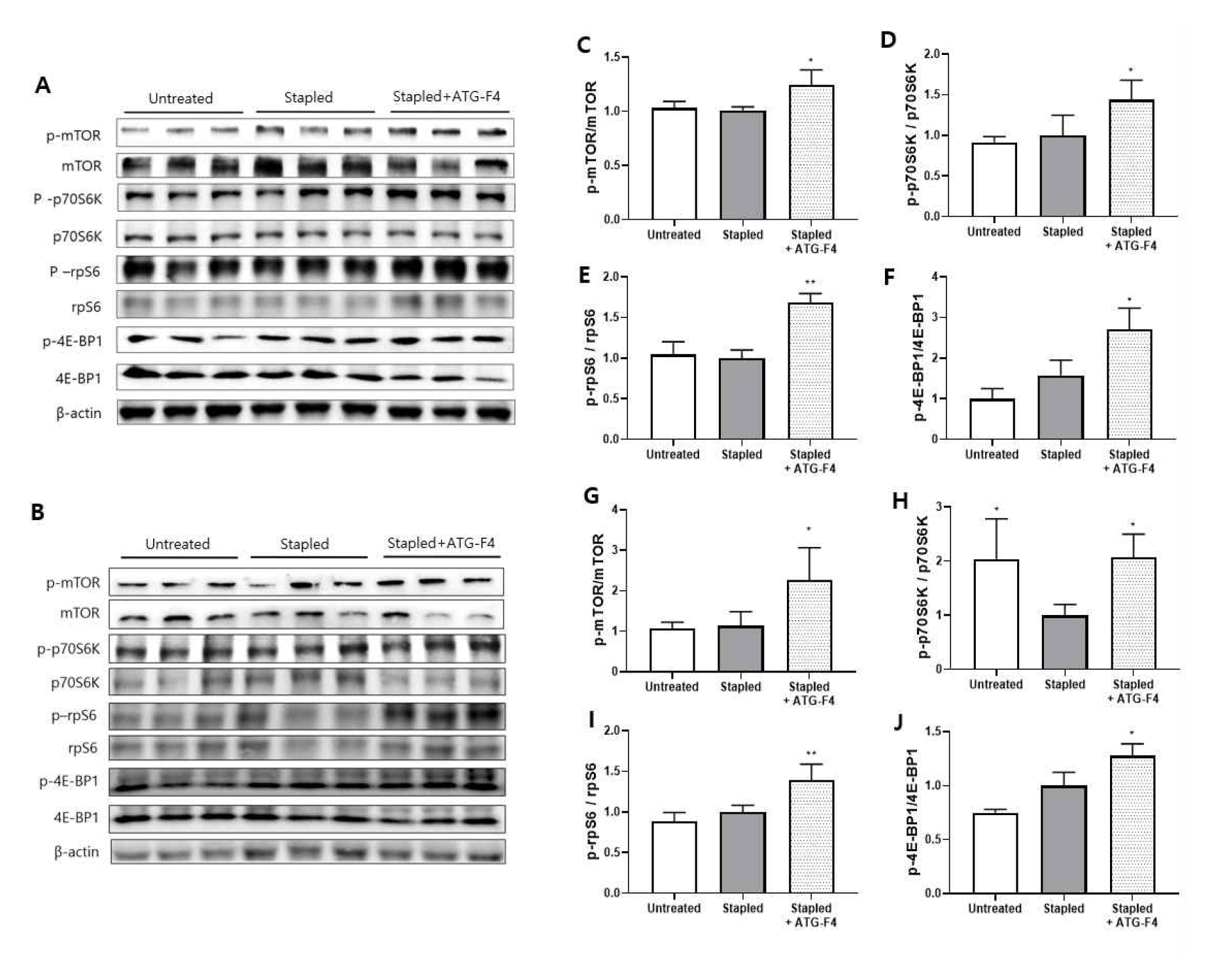
Phosphorylation of muscle synthesis proteins in muscles. (A, B) Representative immunoblotting of the muscle synthesis proteins TA (A) and GA (B), respectively; (C-F) Relative phosphorylation levels (or expression levels) of mTOR (C), and p70S6K (D), rpS6 (E), 4E-BP1 (F) in TA muscle; (G-J) Relative phosphorylation levels (or expression levels) of mTOR (G), and p70S6K (H), rpS6 (I), 4E-BP1 (J) in GA muscle; TA: tibialis anterior; GA: gastrocnemius. Untreated: unstapled group; Stapled: hindlimb immobilization group; Stapled + ATG-F4: hindlimb immobilization + *L. reuteri* ATG-F4 (4.0×10^9^ CFU/day) treated group. Data are presented as means ± SEM *p < 0.05, **p < 0.01 compared to Stapled.

### 3.7 Fecal bacterial community analysis

The changes in gut microbial community were investigated by analyzing fecal samples from each experimental group. Taxonomic abundance analysis revealed a significant increase in the population of Bacteroidetes and a significant decrease in the population of Firmicutes in the ATG-F4 treated group. At the family level, the relative abundance of Muribaculaceae (phylum Bacteroidetes) showed a significant increase (Figure 7A), while the relative abundance of Lachnospiraceae (phylum Firmicutes) and Lactobacillaceae (phylum Firmicutes) showed a significant decrease in the ATG-F4 group, similar to the untreated group (Figure 7B and C).

**Figure 7.**
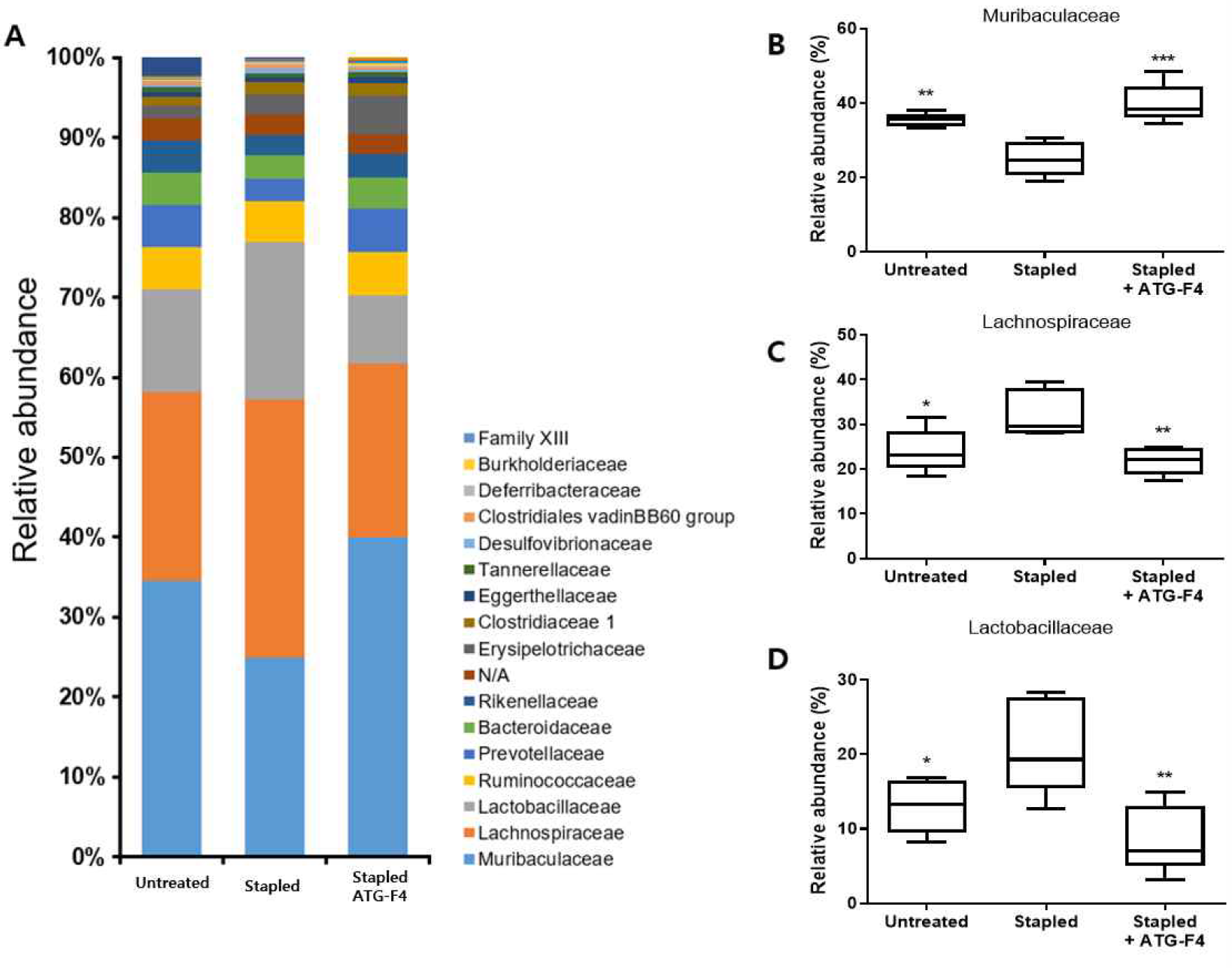
Changes in fecal bacterial community. (A) Stacked bar plot of top 17 abundant family level taxa; (B-D) Relative abundance of Muribaculaceae (B), Lachonospiraceae (C), Lactobacillaceae (D). Untreated: unstapled group; Stapled: hindlimb immobilization group; Stapled + ATG-F4: hindlimb immobilization + *L. reuteri* ATG-F4 (4.0×10^9^ CFU/day) treated group. Data are presented as means ± SEM *p < 0.05, **p < 0.01 compared to Stapled.

### 3.8 Metabolome analysis

The gut microbiome-related metabolites, short-chain fatty acids (SCFAs), were analyzed in serum using LC-MS/MS (Figure 8). The level of butyric acid was increased in the stapled group compared to the untreated group, and the ATG-F4 group showed a greater increase than the stapled group (Figure 8A). The level of acetic acid was not significantly different between the untreated and stapled groups, but the ATG-F4 group showed a higher level of acetic acid than the stapled group (Figure 8B). The levels of propionic acid did not show significant differences among all groups (Figure 8C).

**Figure 8.**
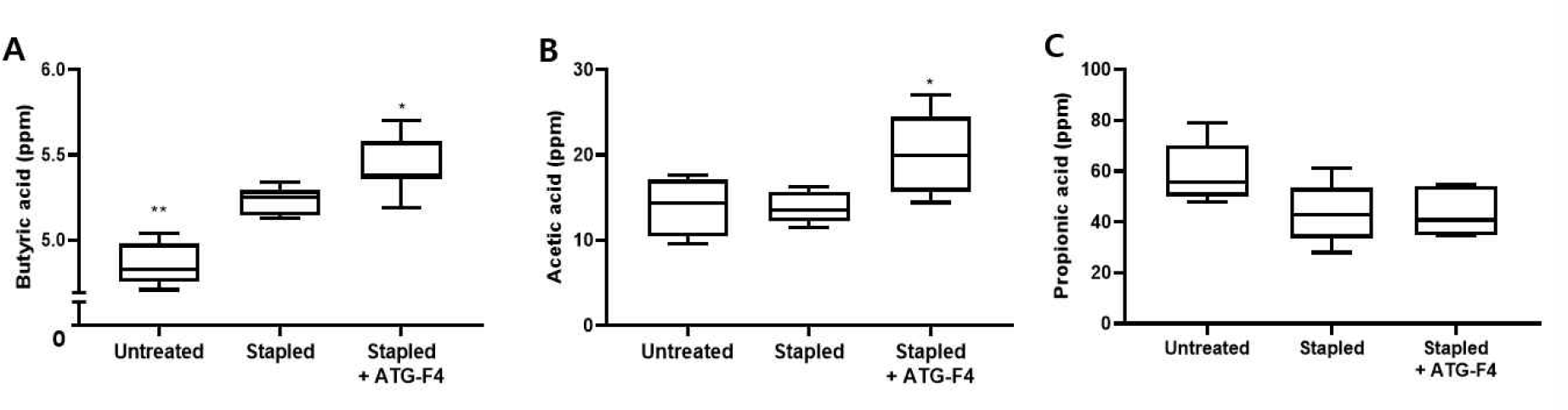
Changes of SCFAs in serum. (A) Butyric acid; (B) Acetic acid; (C) Propionic acid; Untreated: unstapled group; Stapled: hindlimb immobilization group; Stapled + ATG-F4: hindlimb immobilization + *L. reuteri* ATG-F4 (4.0×10^9^ CFU/day) treated group. Data are presented as means ± SEM *p < 0.05, **p < 0.01 compared to Stapled.

## 4 Discussion

The present study aimed to investigate the potential of Lactobacillus reuteri ATG-F4 to ameliorate muscle atrophy and dysfunction. Additionally, we sought to elucidate the underlying mechanism of action in mice with muscle atrophy induced by staple immobilization. Our findings demonstrate that Lactobacillus reuteri ATG-F4 increased muscle mass and improved grip strength and endurance. Moreover, the probiotic was found to suppress inflammation induced by disuse of muscle, indicating its potential to prevent or delay muscle loss. Based on these results, our study suggests that Lactobacillus reuteri ATG-F4 may be a promising candidate for preventing or delaying muscle atrophy by regulating inflammation and improving muscle function.

Increased inflammation levels in inactive muscles are thought to play a significant role in inducing muscle atrophy [25]. The regulation of muscle mass is a complex interplay between catabolic and anabolic processes, where downstream effectors of mTOR, including rpS6 kinase and eIF4E-binding protein (4E-BP1) [26, 27], contribute to protein synthesis and muscle hypertrophy, while muscle atrophy-related genes MuRF1 and Atrogin-1 are responsible for protein breakdown and muscle wasting. It has been suggested that these two pathways are influenced by cytokines and modulated by the presence of inflammation[28]. Additionally, muscle fiber size is an important metric in determining muscle mass and strength, critical for assessing the extent of muscle loss [29-32].

In the previous experiment, no increase in muscle mass was observed in normal animals after 15 days of L. reuteri ATG-F4 administration (S. Figure 1). Similarly, it is speculated that L. reuteri ATG-F4 administration for 2 weeks before staple treatment in the current experiment did not affect muscle mass itself. Moreover, there was no significant difference in muscle recovery between four and seven days after staple removal with L. reuteri ATG-F4 administration (S. Figure 2). Thus, it is expected that L. reuteri ATG-F4 may have suppressed muscle atrophy during the staple period.

In this study, L. reuteri ATG-F4 was analyzed in samples dissected on the fourth day after staple removal. The results showed that L. reuteri ATG-F4 increased muscle fiber size, improved muscle function, and decreased inflammation levels. The increase in muscle fiber size by L. reuteri ATG-F4 strongly supports muscle mass and strength gains [33]. On the other hand, while ATG-F4 increased treadmill endurance, there was no change in soleus muscle mass related to endurance. The increase in endurance due to the administration of L. reuteri ATG- F4 is expected to be associated with the increase in blood levels of BA and AA [34, 35].

The results also showed that L. reuteri ATG-F4 increases phosphorylation of signaling pathway associated with mTOR in muscles and decreases the expression levels of MuRF1. On the other hand, no significant differences were observed in signaling pathway related to muscle synthesis and atrophy between the untreated group and the stapled group. Several papers have shown that signaling pathway related to muscle mass may increase or decrease during treadmill exercise depending on the experimental conditions [36-40]. Studies conducted under experimental conditions similar to the present study, including treadmill exercise and time of dissection, have reported an increase in MuRF1 and mTOR associated with muscle atrophy after treadmill exercise [41, 42]. Therefore, signaling pathway related to muscle atrophy and synthesis after treadmill exercise were increased in both the untreated and stapled group, suggesting that there may be no difference between the two groups. Nevertheless, it is interesting to note that ATG-F4, unlike the other groups, had a positive effect on muscle mass gain associated with muscle atrophy and synthesis.

The imbalance of microbial communities in the gut can cause intestinal barrier function, host metabolism, and signaling pathways to be remodeled. These changes can be directly or indirectly related to gut dysbiosis and inflammation [43, 44]. In addition, inflammation occurring within the body, such as that caused by lung or breast cancer, can lead to changes in gut microbiota [45]. It is believed that changes in gut microbiota in the staple immobilization model occur because inflammation originating from the muscles may have affected the gut.

Administration of L. reuteri ATG-F4 resulted in changes in gut microbiota; however, the Lactobacillus cluster did not increase with ATG-F4 administration (S. Figure 3A). Treatment with L. reuteri ATG-F4 led to an increase in Muribaculaceae and a decrease in Lachnospiraceae and Lactobacillaceae, which tended to match the normal control group without stapled.

The abundance of Muribaculaceae is mostly decreased in many inflammation-related diseases [46-48]. However, Lachnospiraceae and Lactobacillaceae have been reported to increase or decrease depending on the inflammation model [46, 49-51]. Regarding changes in microbial balance within the gut, more studies are needed to investigate whether L. reuteri ATG-F4 can alter the composition ratio of Muribaculaceae, Lachnospiraceae, and Lactobacillaceae. However, it is noteworthy that L. reuteri ATG-F4 induced changes in the gut microbiota composition similar to those observed in healthy animals. Therefore, administering L. reuteri ATG-F4 could be a potential probiotic for improving gut microbiota imbalance.

The previously mentioned Muribaculaceae, Lachnospiraceae, and Lactobacillaceae are dominant strains that account for approximately 70% of the total gut microbiota. These strains have the ability to ferment complex carbohydrates and dietary fiber, which leads to the production of short-chain fatty acids (SCFAs) as a byproduct [52]. SCFAs are essential for maintaining gut function, as they serve as an energy source for colon cells, reduce inflammation, and improve intestinal motility [53-55]. Additionally, evidence is increasingly accumulating to suggest that SCFAs have beneficial effects on skeletal muscle metabolism and function [13, 56].

BA, one of the SCFAs, has been reported to have anti-inflammatory properties and may act as an energy source for muscle [57-59]. It has also been reported that administration of BA to an aged rat model increases muscle mass and fiber size, and the mechanism underlying the positive effects on muscle involves increased mTOR and inhibition of MuRF1[60, 61]. In addition, AA may benefit muscle function by improving glycogen replenishment and increasing acetyl-CoA concentrations in skeletal muscle [62, 63]. Due to the effects mentioned above, BA and AA are considered attractive SCFA metabolites for inhibiting muscle atrophy and increasing endurance [64].

The increase in blood concentrations of BA and AA following L. reuteri ATG-F4 administration suggests that L. reuteri ATG-F4 may stimulate the growth and activity of indigenous bacteria. These findings suggest that L. reuteri ATG-F4 can increase muscle mass in atrophied muscles, which may be accompanied by BA-induced upregulated mTOR and downregulated MuRF1 pathways; and the increase in AA may have contributed to the improvement of muscle endurance. However, unlike the similar bacterial community ratios observed in NC and L. reuteri ATG-F4, BA and AA in serum were increased only in L. reuteri ATG-F4. Comparing the relative abundance of bacterial communities at the genus level with the concentration Increase/decrease pattern of BA and AA, a similar pattern was observed with Parasutterella, Monoglobus, Faecalibaculum (S.Figure 3B-D). However, since these strains have a very low relative proportion, further experiments are needed to discuss the increase of BA and AA by ATG-F4 administration.

Many studies have shown that whole body inflammation interacts with the gut, and that gut immune function can modulate whole body inflammation [65, 66]. In addition, gut-derived substances produced in the gut can be transported throughout the body and exert various functions [67]. As discussed above, L. reuteri ATG-F4 appears to improve muscle mass and function by restoring gut microbial communities, increasing serum BA and AA levels, and suppressing inflammation. Based on our findings, L. reuteri ATG-F4 may regulate the gut microbiota-SCFAs (BA, AA)-muscle axis, thereby ameliorating muscle atrophy.

## 5 Conclusion

In this study, the novel composition ATG-F4 was found to significantly increase muscle mass and fiber size, while improving muscle strength and exercise performance in a model of immobilized muscle atrophy. The mechanism of action was attributed to the activation of a series of mTOR pathways related to muscle synthesis and the inhibition of muscular atrophy factors, which was accompanied by alterations in the relative composition of the intestinal microbiota and anti-inflammatory effects. These findings suggest that ATG-F4 has potential as a prophylactic or therapeutic agent for muscle atrophy.

## 6 Author contributions

DY L designed and conducted animal experiments and analysed data and wrote the manuscript. YS L carried out the animal experiment and edited the manuscript. GS P and SH K analyzed meta-analyses data of fecal microbiota and participated in manuscript writing. JY L and YK L participated in analysis of SCFAs. DY J and YH L gave appropriate advices from preliminary data of staple model. JH K: supervised the manuscript. All coauthors contributed to the manuscript, and have approved the final manuscript.

## 7 Data availability

The datasets generated and/or analyzed during the current study are available at NCBI’s repository. The raw sequence data of bacterial community sequencing are submitted to NCBI SRA database (NCBI BioProject PRJNA694467, https://www.ncbi.nlm.nih.gov/bioproject/694467).

## 8 Acknowledgments

The present study was financially supported by Ministry of SMEs and Startups (grant no. S2912175; Development of functional health food materials to prevent sarcopenia using probiotics). We gratefully acknowledge the assistance of MSS.

---

**S. Figure 1.**
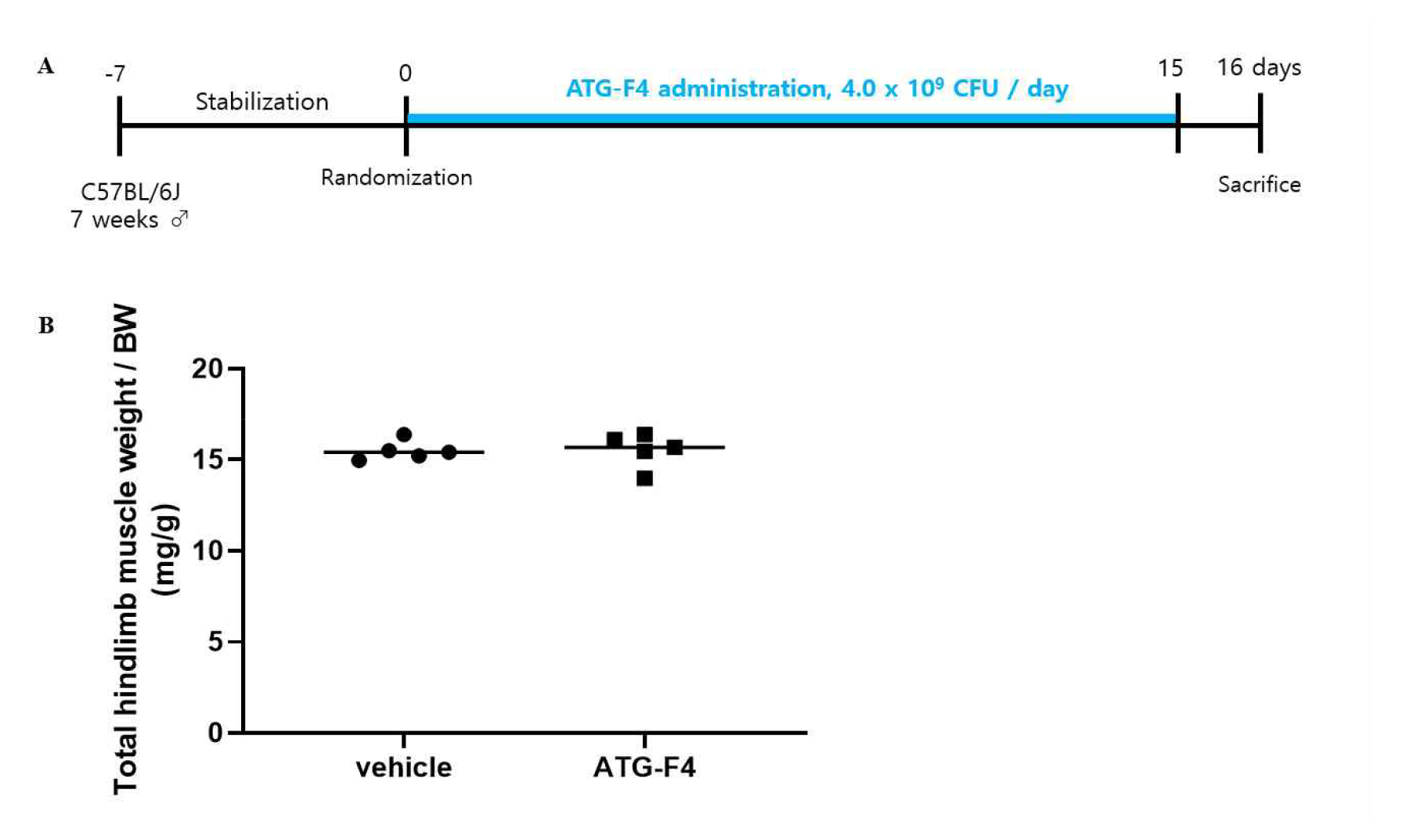
Animal experiment schedule (A) and results of changes in hindlimb muscle mass (B).

**S. Figure 2.**
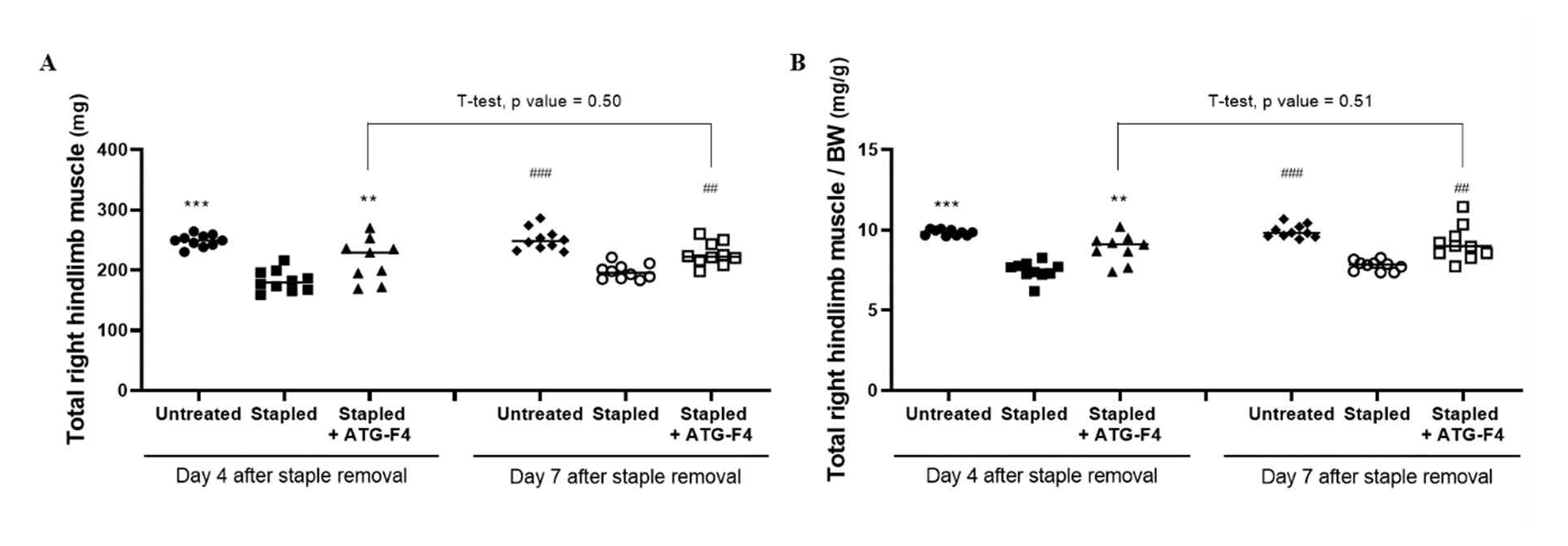
Muscle mass values at 4 and 7 days after staple removal. Absolute values (A), relative values(B). Untreated: unstapled group; Stapled: hindlimb immobilization group; Stapled + ATG-F4: hindlimb immobilization + *L. reuteri* ATG-F4 (4.0×10^9^ CFU/day) treated group. One-way ANOVA Dunnett’s test ***p<0.001, **p<0.01 vs Stapled group in 4days, One-way ANOVA Dunnett’s test ###p<0.001, ##p<0.01 vs Stapled group in 7days, T-test Stapled + ATG-F4 group in 4days vs Stapled + ATG-F4 group in 7days

**S. Figure 3.**
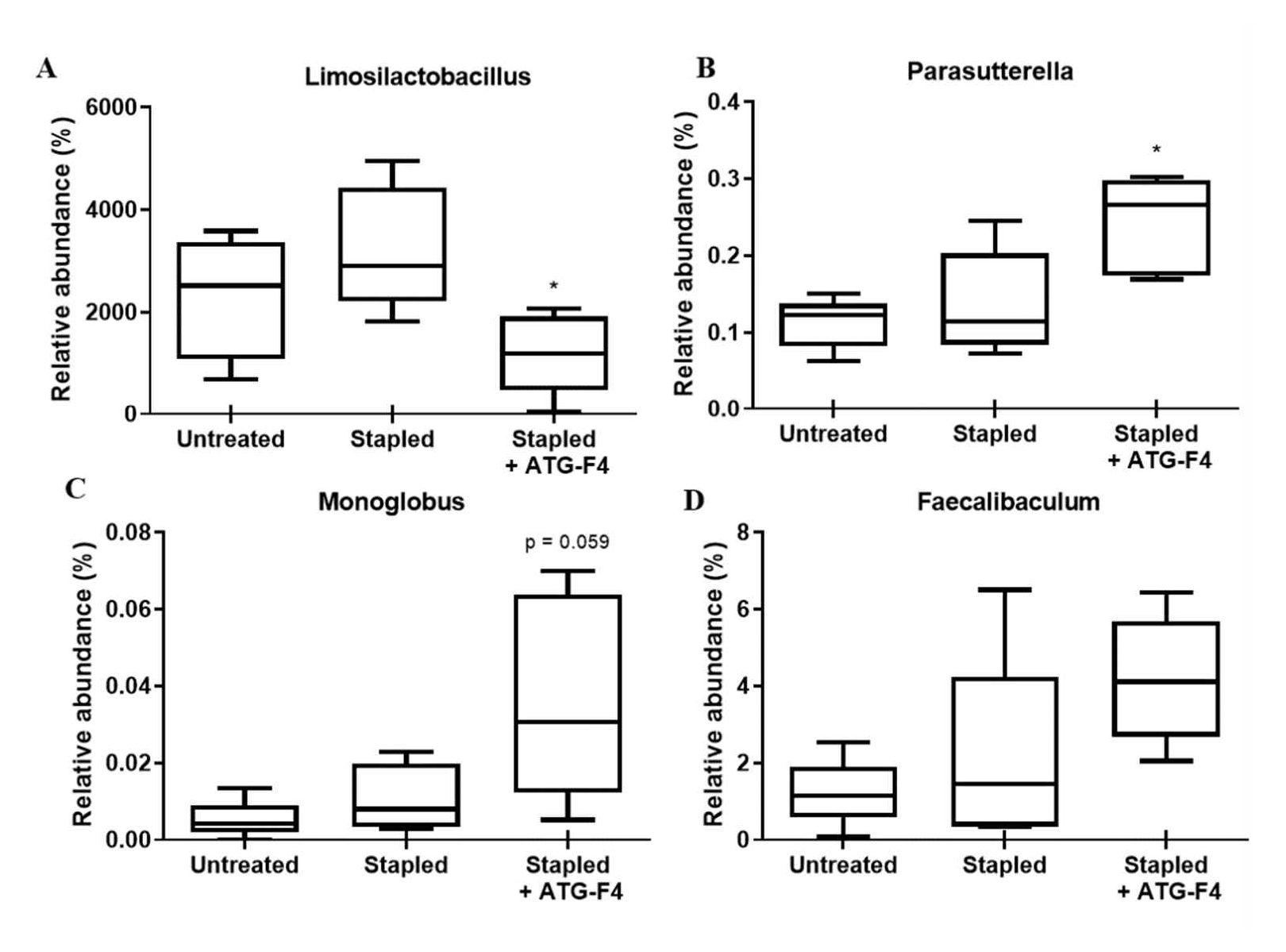
Changes in fecal bacterial communities at the Genus level. Limosilactobacillus (A), Parasutterella (B), Monoglobus (C), Faecalibaculum (D), Untreated: unstapled group; Stapled: hindlimb immobilization group; Stapled + ATG-F4: hindlimb immobilization + *L. reuteri* ATG-F4 (4.0×10^9^ CFU/day) treated group.

